# High contiguity *Arabidopsis thaliana* genome assembly with a single nanopore flow cell

**DOI:** 10.1101/149997

**Authors:** Todd P. Michael, Florian Jupe, Felix Bemm, S. Timothy Motley, Justin P. Sandoval, Olivier Loudet, Detlef Weigel, Joseph R. Ecker

## Abstract

While many evolutionary questions can be answered by short read re-sequencing, presence/absence polymorphisms of genes and/or transposons have been largely ignored in large-scale intraspecific evolutionary studies. To enable the rigorous analysis of such variants, multiple high quality and contiguous genome assemblies are essential. Similarly, while genome assemblies based on short reads have made genomics accessible for non-reference species, these assemblies have limitations due to low contiguity. Long-read sequencers and long-read technologies have ushered in a new era of genome sequencing where the lengths of reads exceed those of most repeats. However, because these technologies are not only costly, but also time and compute intensive, it has been unclear how scalable they are. Here we demonstrate a fast and cost effective reference assembly for an *Arabidopsis thaliana* accession using the USB-sized Oxford Nanopore MinION sequencer and typical consumer computing hardware (4 Cores, 16Gb RAM). We assemble the accession KBS-Mac-74 into 62 contigs with an N50 length of 12.3 Mb covering 100% (119 Mb) of the non-repetitive genome. We demonstrate that the polished KBS-Mac-74 assembly is highly contiguous with BioNano optical genome maps, and of high per-base quality against a likewise polished Pacific Biosciences long-read assembly. The approach we implemented took a total of four days at a cost of less than 1,000 USD for sequencing consumables including instrument depreciation.

## Introduction

The first *Arabidopsis thaliana* genome assembly, from the Columbia (Col-0) accession, was released 17 years ago, based on Sanger sequencing of a BAC tiling path [1]. Subsequently this reference assembly has been continuously improved to a gold standard, resulting in the current TAIR10 version with only 29 non-centromeric mis-assemblies [2] and 92 gaps with unknown bases (‘Ns’). TAIR10 is also missing about 25 Mb of repeat (mostly centromere) sequence [3]. The advent of high density microarrays and later Illumina short-read sequencing led to several efforts to access the nucleotide and structural diversity of genomes from other *A. thaliana* accessions [3–7]. Most of these efforts were reference based, revealing only limited structural changes. Various attempts at reference guided or *de novo* assembly with short reads have been pursued [8,9], but these also suffered from being unable to accurately reconstruct larger insertions as well as presence/absence polymorphisms of medium-copy repeats.

More recently third generation sequencing technologies, such as SMRT Sequencing from Pacific Biosciences (PB) have made it possible to assemble highly contiguous genomes [10,11] since the read lengths exceed those of the major repeats, although highly repetitive regions such as the ribosomal DNA (rDNA) and centromeres still remain unassembled with third generation technologies due to limitations in read length and error rate. Oxford Nanopore (ONT) MinION sequencing has overcome at least one of these challenges by producing reads that exceed 200 kb, but still have 5-15% error (vR7.3; [12]). Optical genome mapping, on the other hand, allows long-range scaffolding and identification of large structural genome variation, however only in combination with long-read sequencing, and it is not able to resolve the underlying genome sequence independently [2].

In this effort we have generated a highly contiguous *A. thaliana* genome assembly from 1 μg genomic DNA on just a single ONT MinION flow cell (R9.4). We validated the contiguity and the quality with BioNano optical maps and a second assembly generated from long reads produced on a PB RSII platform. The final assembly is on par with that of the current gold standard Arabidopsis TAIR10 assembly, and was generated in four days for under 1,000 USD for sequencing consumables and instrument depreciation.

## Results

We extracted high-molecular weight DNA from leaf tissue of the *A. thaliana* accession KBS-Mac-74 (ecotype 1741), and used 1 μg to prepare a one dimensional (1D) ONT sequencing library (SQK-LSK108). This DNA was sequenced using one (1) ONT MinION R9.v4 flowcell for 48 hrs, resulting in 300,053 fast5 files (sequencing reads), which were subsequently analyzed using the albacore recurrent neural network (RNN) base-caller (v0.8.4). The average length was 11.4 kb (N50 7.5 kb), a total of 3.4 Gb of base-called sequence (Figure 1). There were four (4) reads greater than 200 kb, including one of 269 kb, 14 greater than 100 kb, and 2,317 greater than 50 kb. The reads had an average quality of Q7.3. Plotting GC content against read length revealed that some of the longest reads were skewed toward low GC (25%) compared to the average in the *A. thaliana* reference genome (36%). As a comparison we generated 8.4 Gb of Pacific Biosciences RSII (PB) long reads from the same accession, and BioNano Genomics optical genome maps to confirm and validate the assemblies (Methods).

**Figure 1.**
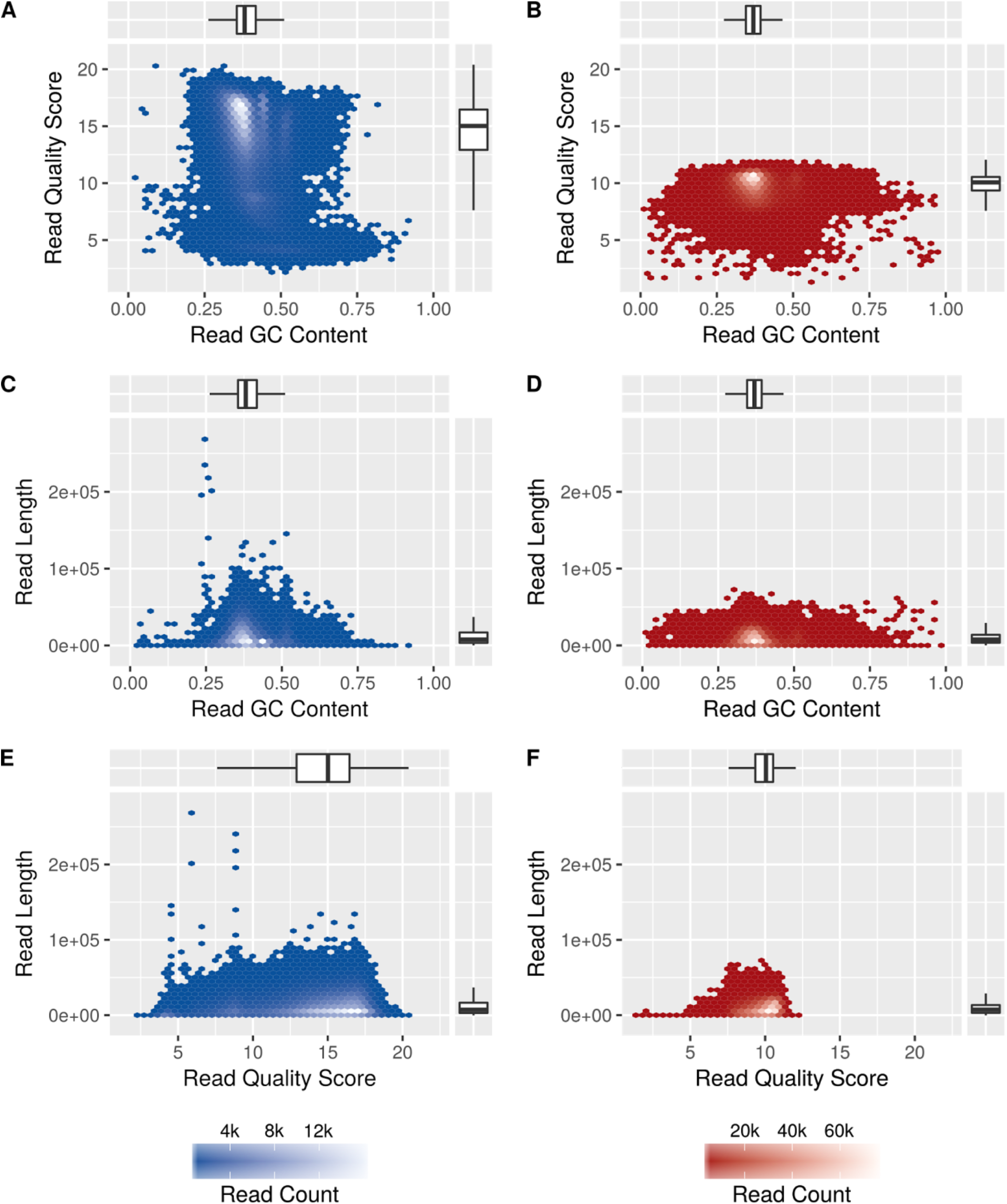
*Arabidopsis thaliana* Oxford Nanopore (ONT) sequence compared to PacBio (PB) sequence. (A-B) ONT (blue) and PB (red) read quality versus GC content. (C-D) Read length versus GC content. (E-F) Read length (bp) versus quality score.

While several long-read, read error-tolerant assemblers produce high quality genomes, such as Canu [Citation error] and Falcon [13], they are compute intensive for larger eukaryotic genomes. In contrast, minimap/miniasm is a new assembler specifically designed to handle long error-prone reads in a fraction of the time [14]. We leveraged both assemblers to assemble raw ONT and PacBio reads (fastq). All four initial assemblies compared similar in common assembly metrics and showed a high contiguity (Table 1). Of note, the ONT minimap/miniasm (ONTmin) assembly has the least contigs (62) and the smallest assembled genome size (110.9 Mb), but the second longest N50 (11.5 Mb) and highest maximal contig size (13.8 Mb). The PacBio Canu (PBcan) assembly was superior in all other metrics.

**Table 1.** Metrics of the KBS-Mac-74 Oxford Nanopore (ONT) and Pacific Biosciences (PB) assemblies.

| **Name** | **Iteration (#)** | **Contig (#)** | **Total length (bp)** | **N50 length (bp)** | **Longest (bp)** |
| --- | --- | --- | --- | --- | --- |
| ONTcan | 0 | 82 | 113,402,589 | 8,491,723 | 13,170,991 |
| ONTcan | 1 | 82 | 115,632,969 | 8,666,702 | 13,418,669 |
| ONTcan | 2 | 82 | 115,735,933 | 8,671,757 | 13,431,826 |
| ONTcan | 3 | 82 | 115,781,606 | 8,673,416 | 13,434,963 |
| ONTcan | 4 | 82 | 117,857,778 | 8,825,005 | 13,671,933 |
| PBcan | 0 | 119 | 122,797,851 | 13,362,607 | 16,203,764 |
| PBcan | 1 | 116 | 122,817,504 | 13,378,182 | 16,222,950 |
| PBcan | 2 | 113 | 122,746,321 | 13,380,370 | 16,226,171 |
| PBcan | 3 | 111 | 122,681,645 | 13,381,241 | 16,226,984 |
| PBcan | 4 | 111 | 122,486,187 | 13,361,814 | 16,200,459 |
| ONTmin | 0 | 62 | 110,880,468 | 11,450,026 | 13,809,770 |
| ONTmin | 1 | 62 | 117,109,669 | 12,091,692 | 14,572,214 |
| ONTmin | 2 | 62 | 117,377,831 | 12,118,333 | 14,600,216 |
| ONTmin | 3 | 62 | 117,441,618 | 12,123,124 | 14,605,848 |
| ONTmin | 4 | 62 | 119,502,799 | 12,334,794 | 14,864,299 |
| PBmin | 0 | 114 | 124,752,783 | 6,618,840 | 13,276,998 |
| PBmin | 1 | 114 | 120,562,781 | 6,390,883 | 12,830,360 |
| PBmin | 2 | 114 | 120,313,229 | 6,377,043 | 12,801,432 |
| PBmin | 3 | 114 | 120,275,462 | 6,375,227 | 12,798,469 |
| PBmin | 4 | 114 | 120,047,081 | 6,362,171 | 12,770,538 |

The minimap/miniasm assembly strategy does not include an initial error correction of the raw reads such as implemented in the standard Canu pipeline. Therefore, the minimap/miniasm assembly has many local mis-assemblies compared to the Canu assemblies (Table 2). We polished the assemblies using three (3) iterations of racon [15], followed by one (1) round of polishing using Illumina PCR-free paired-end reads with pilon [16] (Table 2). In general, racon significantly improved the ONT assemblies for total length (Figure 2A), and increased the ONTmin N50 length and longest contig to 12.3 and 14.8 Mb respectively. The minimap/miniasm assemblies had a high local assembly problem as determined by split read mapping (Table S1) being an order of magnitude higher than the raw Canu assemblies (Figure 2B). In addition, all of the assemblies benefited from several rounds of racon for overall variants as well as SNPs, with the exception of the PBcan assembly (Figure 2C,D). Overall, after three rounds of racon and one round of pilon the assemblies were similarly low for variants (Figure 2; Table 2).

**Figure 2.**
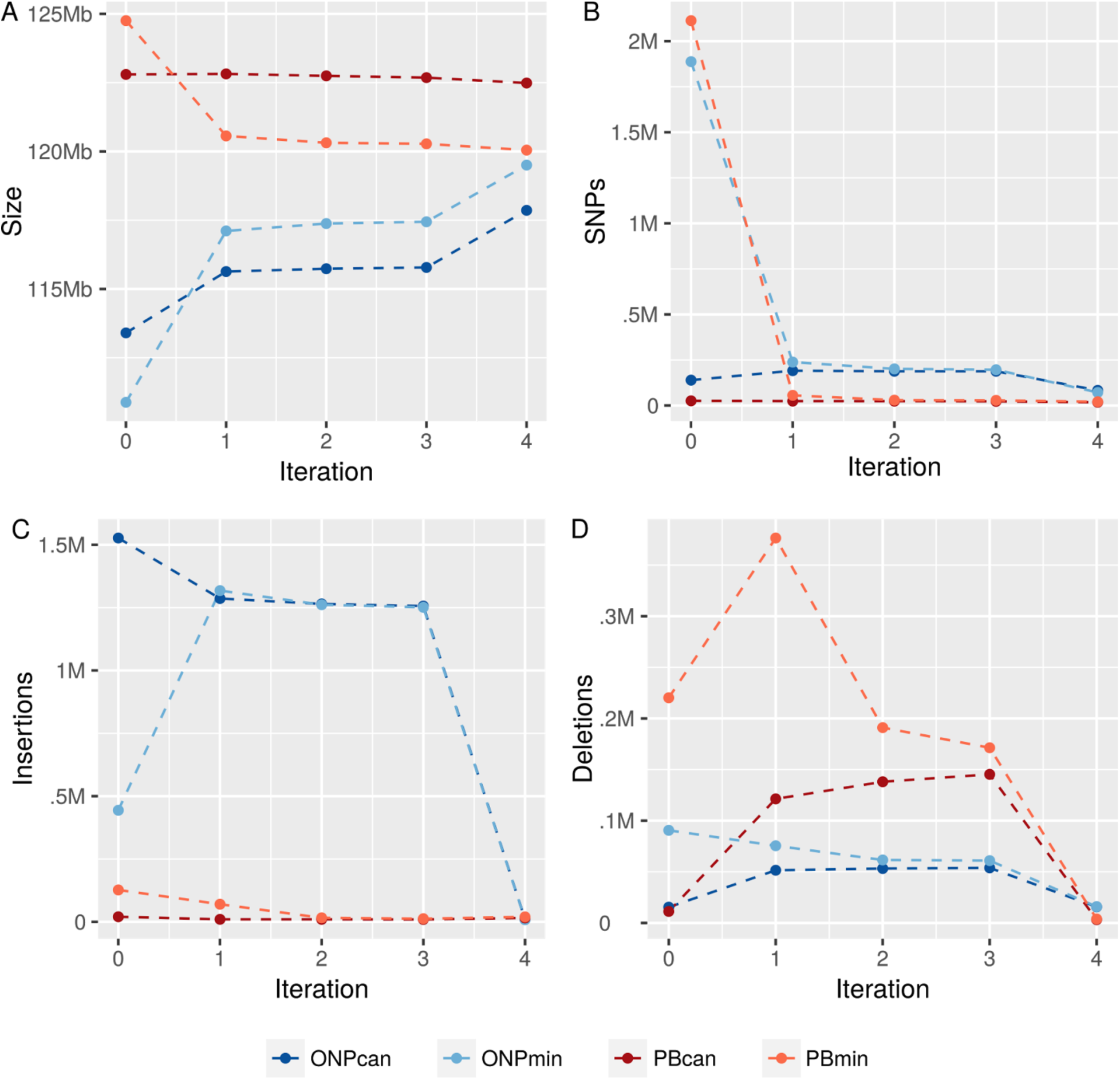
Oxford Nanopore (ONT) and PacBio (PB) assembly comparisons. (A) Final assembled genome size versus assembly iterations. (B) SNPs versus iterations. (C) Insertions versus iterations. (D) Deletions versus iterations. Assembly types: Canu, can; Minimap/miniasm, min. Assembly iterations: 0, raw assembly; 1, racon 1x; 2, racon 2x; 3, racon 3x; 4, pilon 1x.

**Table 2.** Variant mapping metrics of Illumina PCR-free paired-end reads against the assemblies.

| **Name** | **Iteration**<br>**(#)** | **Total Variants**<br>**(#)** | **SNPs**<br>**(#)** | **Insertions**<br>**(#)** | **Deletions**<br>**(#)** |
| --- | --- | --- | --- | --- | --- |
| ONTcan | 0 | 1,681,854 | 139,358 | 1,526,965 | 15,393 |
| ONTcan | 1 | 1,529,269 | 191,313 | 1,286,344 | 51,513 |
| ONTcan | 2 | 1,506,201 | 188,078 | 1,264,824 | 53,208 |
| ONTcan | 3 | 1,498,396 | 188,033 | 1,256,415 | 53,852 |
| ONTcan | 4 | 108,830 | 83,303 | 9,816 | 15,705 |
| PBcan | 0 | 57,430 | 25,842 | 20,343 | 11,237 |
| PBcan | 1 | 155,751 | 24,291 | 10,122 | 121,329 |
| PBcan | 2 | 172,047 | 23,978 | 10,016 | 138,044 |
| PBcan | 3 | 177,711 | 22,862 | 9,508 | 145,332 |
| PBcan | 4 | 36,726 | 17,792 | 15,754 | 3,175 |
| ONTmin | 0 | 2,422,153 | 1,887,582 | 443,925 | 90,587 |
| ONTmin | 1 | 1,631,836 | 238,425 | 1,317,831 | 75,486 |
| ONTmin | 2 | 1,524,650 | 201,304 | 1,261,647 | 61,596 |
| ONTmin | 3 | 1,508,869 | 196,529 | 1,251,302 | 60,951 |
| ONTmin | 4 | 96,353 | 71,724 | 8,713 | 15,911 |
| PBmin | 0 | 2,459,812 | 2,112,559 | 127,003 | 220,235 |
| PBmin | 1 | 502,726 | 56,016 | 70,240 | 376,453 |
| PBmin | 2 | 236,765 | 29,699 | 16,055 | 191,006 |
| PBmin | 3 | 212,493 | 28,536 | 12,629 | 171,323 |
| PBmin | 4 | 44,630 | 20,913 | 19,850 | 3,865 |

BioNano Genomics (BioNano) Irys maps have been very useful in identifying mis joins and analyzing the contiguity of short- and long-read assemblies [2,17,18]. We generated 265 optical genome maps with lengths up to 4.8 Mb and an N50 of 695 kb, and assessed the quality of the assemblies by screening for chimeric contigs and mis-assemblies such as collapsed and artificially expanded regions (Table S2). While we found one single chimeric contig in ONTcan, all other approaches assembled this region perfectly (Figure 3A,B). Artificially expanded regions occur only in the miniasm assemblies, with the exception of four such regions in the ONTcan raw assembly. These expanded regions disappear after the first racon round. Within the final ONTmin contigs we identify four contracted sequences within highly repetitive regions, totalling 128,258 bp (one example in Figure 3C). One example of an erroneously extended region of ~18 kb, most likely a repeat, within ONTmin is shown in Figure 3D.

**Figure 3.**
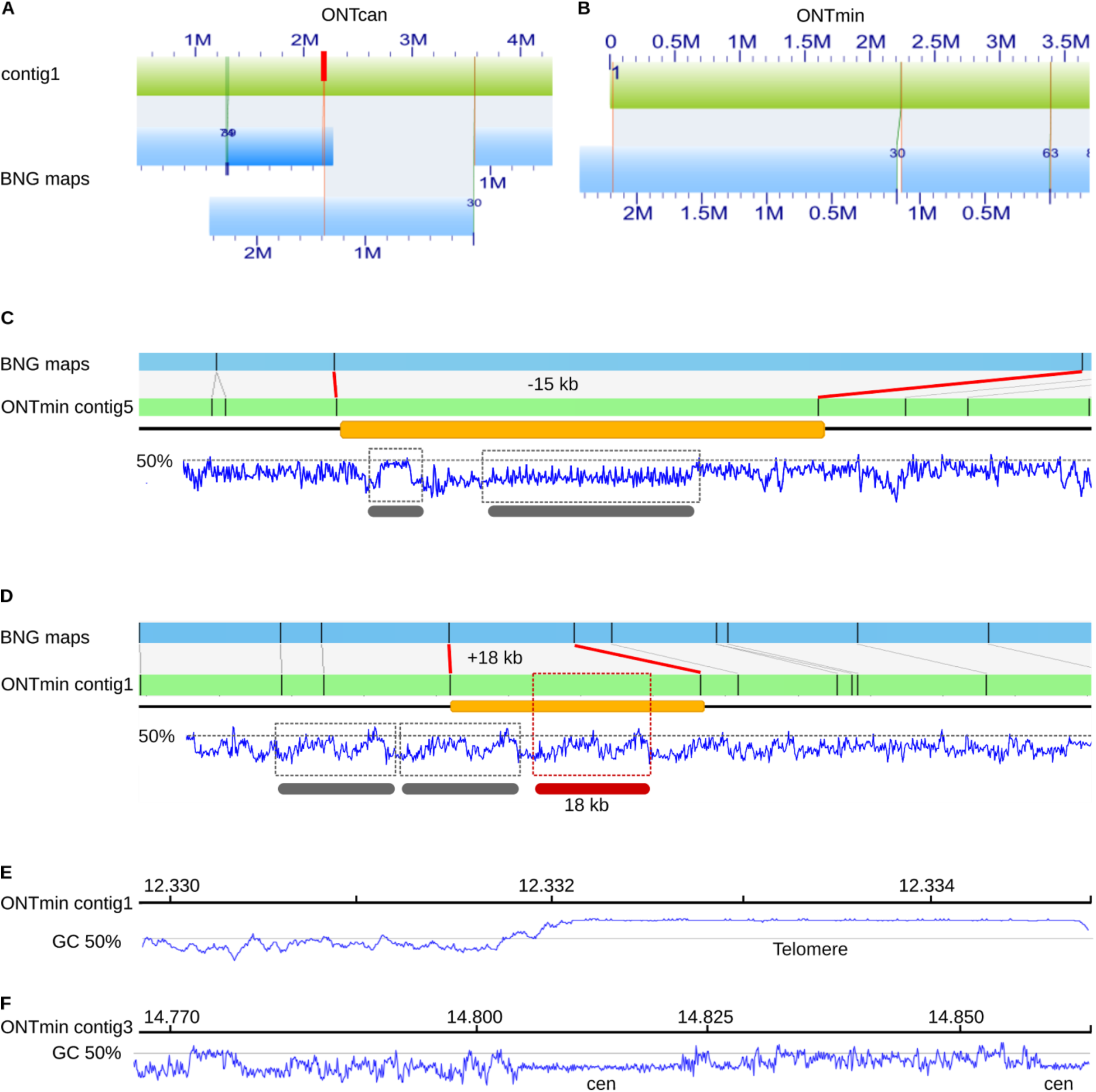
BioNano Genomics (BNG) maps identify mis-assemblies and hard to assemble regions. BNG cmap_30 (Blue) identified (A) a chimeric ONTcanu contig1 (green) and (B) the correct assembled contig1 in the ONTmin assembly (green). The chimeric position is indicated with a red bar. We highlight (C) a collapsed region in ONTmin contig5 in which approximately 15 kb sequence are missing from one of two potential repeat regions as identified by the GC pattern (grey bars). In contrast, (D) shows an artificially expanded region of approximately 18 kb, with the duplicated repeat region highlighted (red bar, 18 kb). (E) ONTmin assembly resolves various telomere regions for example after 12.332 Mb on contig1, as outlined by a GC plot (blue line). (F) ONTmin also resolves short centromere arrays as shown towards the end of contig3 (blue, GC plot).

We further measured global base quality through *in silico* labeling of the contigs at Nt.BspQI sites and comparing to BioNano maps, also based on Nt.BspQI restriction sites. This analysis also revealed false-positive (FP) and false-negative (FN) Nt.BspQI sites. We compared the five iterations per assembly against the KBS-Mac-74 BioNano maps (Table S2). The raw ONTmin assembly (iteration: 0) failed to produce good alignments with optical maps due to high nucleotide errors, causing little overlapping labels. The first polishing step enabled testing of 115.9 Mb, which increased to 118.4 Mb in the final ONTmin assembly (iteration: 4). FN/FP ratios decreased from 0.33/0.08 to 0.02/0.04, with a dramatic shift due to the final pilon polish (iteration: 4). While changes were marginal between racon iteration 2 and 3, the pilon polishing step improves all assembly types, decreasing the FN/FP ratios in ONTcan (iteration: 4) from 0.12/0.33 to 0.04/0.01 (Table S2, Figure 2).

Minimap/miniasm does not attempt to assemble repeat sequences [14], so we determined the difference in the amount of repeats between the four assemblies. While we were not able to identify large repeats, such as the nucleolar organizer rDNA at the short arms of chromosome 2 and 4 in our assemblies, we can see telomeric and centromeric repeats, that were most likely part of a single read. Examples include a 2.8 kb telomere repeat on contig1 (Figure 3E) and two centromeric repeat arrays of 16.5 kb and 4.8 kb at the end of contig3 (Figure 3F). Canu on the other hand assembles larger repeats and engages all rDNA-encoding BioNano maps (Figure 3). The PBcan assembly has an order of magnitude more centromere and rDNA sequence than any of the other assemblies (Table S3). Repeats called using RepeatMasker [19] reveals a similar number of repetitive elements in the assemblies (data not shown). As an additional proxy for the quality of the ONTmin assembly we predicted genes. Similar numbers of genes were predicted for each assembly (data not shown), and while PBcan had the highest identity with Araport11 predicted protein coding genes [20], the ONTmin was very similar in quality (Table S4). Overall, polishing brought the ONT assemblies to a comparable level in base quality and contiguity.

A major motivation for *de novo* assemblies is the identification of functionally important sequences missing from the reference genome assembly. Function can, for example, be assigned by genetic mapping, including Quantitative Trait Locus (QTL) mapping. We therefore analyzed a fine-mapped QTL region [21], where the Bur-0 accession has two copies of the At4g30720 gene, of which Col-0 has only one copy. The two copies are spaced about 10 Mb apart on chromosome 4 in Bur-0, and are defined by two interacting QTL, SG3 and SG3i [21].

We tested the utility of having a rapidly created, highly contiguous reference genome from a non-Col-0 accession for validating this QTL. Blast searches with the Col-0 At4g30720 genomic sequence against our KBS-Mac-74 ONTmin assembly recovered two hits (99.3 and 98.2% identity) 10 Mb apart on the same contig, consistent with the structure identified in Bur-0 (Figure 4). ONTmin was the only assembly where both At4g30720 copies were found on the same contig (data not shown). We then asked if the breakpoints of the SG3i locus could be defined by aligning the SG3i region of our ONTmin assembly to Col-0 (TAIR10). We mapped Col-0 left and right breakpoints at 5,771,424 bp and 5,773,387 bp respectively, spanning 1,963 bp and containing a predicted transposable element with reverse transcriptase homology (At4g08995). The corresponding KBS-Mac-74 region spanned 39,273 bp and indeed contained the QTL gene At4g30720. In this respect, it seems that Bur-0 and KBS-Mac-74 share the same sequence, which is missing from the Col-0 reference accession. More detailed analysis of the KBS-Mac-74 SG3i region revealed that it contained four fragments of the transposable element consistent with several rounds of nested transposition, which partially explains why it was impossible to amplify from our BAC clone using PCR.

**Figure 4.**
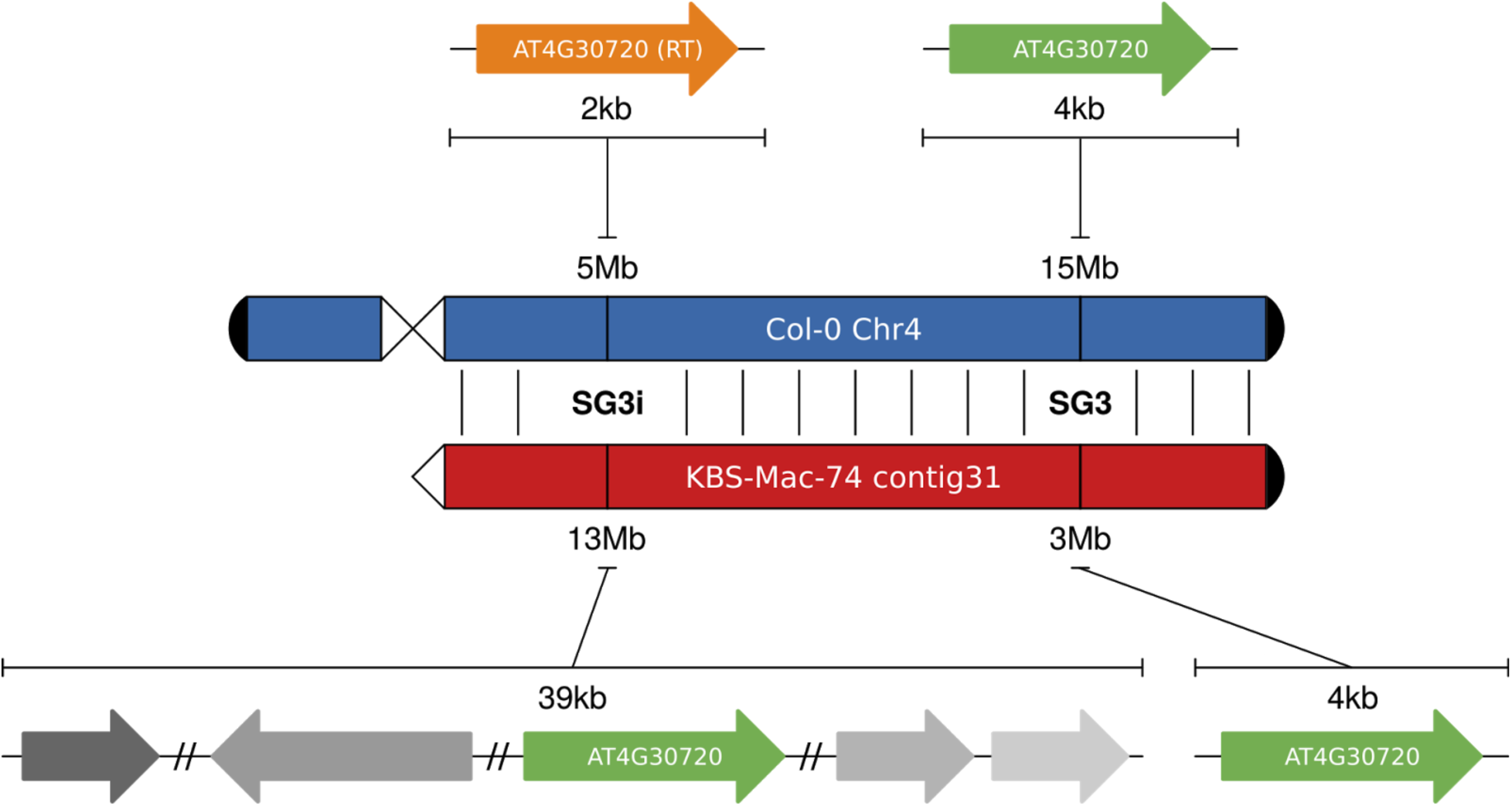
Resolution of the At4g30720 duplication using the KBS-Mac-74 ONTmin genome. Col-0 only has one copy of At4g30720 on the bottom (15 Mb) of chromosome 4 (Chr4). KBS-Mac-74 (ONTmin) assembly has two copies of At4g30720; one at the beginning of contig31 that corresponds to the SG3 QTL location, and one at the bottom of contig31 that overlaps with the SG3 interacting (SG3i) region in the middle of chromosome 4. At the SG3i locus, the Col-0 gene At4g08995 (889 bp), which is annotated as a transposable element (RT; putative reverse transcriptase), is replaced in the KBS-Mac-74 ONTmin assembly with a 39 kb expansion that includes a duplicated copy of At4g30720. Fragments of the TE (different grey arrows) are scattered across the KBS-Mac-74 region consistent with several rounds of transposition resulting in this complex rearranged region.

## Discussion

In one ONT MinION run we sequenced and assembled an *A. thaliana* accession (KBS-Mac-74) to a high continuity and base quality on par with the current “gold standard” TAIR10 assembly of the Col-0 reference accession. The TAIR10 assembly has 94 contigs [1], compared to the 62 contigs of our nanopore assembly, suggesting the underlying 1ONT assembly has higher contiguity. While the initial quality of the minimap/miniasm assembly was lower than Canu, several rounds of racon and one round of pilon produced an assembly on par with PacBio in both contiguity and base quality. Library prep took ~3 hrs, sequencing on the MinION commenced for 48 hrs, assembly with minimap/miniasm less than one hour, consensus with racon (3x) 12 hrs, and polishing with pilon 24 hrs. Together this assembly cost under 1,000 USD in consumables and instrument depreciation and took less than a week of actual time.

The direct comparison of 1D ONT (v9.4) and PacBio (RSII) reads (Figure 1) revealed that 1D ONT reads surpassed the latter in both lengths and quality (Figure 1 E,F). The longest ONT reads (>200 kb) did have a skewed GC content (Figure 1C), but in general the highest quality reads were focused around the core *A. thaliana* GC content of 36% (Figure 1A). In contrast the highest quality PacBio reads spanned the GC content range (Figure 1B). While the new PacBio Sequel platform is generating even longer reads (>60kb, [22]), we routinely generate reads between 200 and 800 kb on the ONT MinION (v9.4). These extremely long reads provide a completely new opportunity to assess highly nested transposable elements and repeat regions that until now were inaccessible to DNA sequencing technologies. Nevertheless, we were unable to use these reads to resolve centromeres and rDNA cassettes because in *A. thaliana* these repeats have high identity (~100%) and span many Mb, which exceeds the quality and length of even our best ONT datasets.

While we have had great success with Canu [11] for PacBio long-read assemblies[11], the assembly of ONT reads took several days (5-10 d depending on the dataset), as compared to the minimap/miniasm raw assemblies, which usually took less than 30 min. The trade-off is that the minimap/miniasm assemblies have substantially lower per-base quality than the canu assemblies and don’t assemble highly repetitive regions such as centromeres and rDNA (Figure 2; Table S3). Minimap/miniasm takes a “correction-free” assembly approach [14], which works particularly well with exceptionally long ONT, error-prone reads. Not only did several rounds of racon improve the SNP, deletion and insertion rate it also properly extended the ONT assemblies to 100% of the non-repetitive genome size (Figure 2A). Most importantly, a final pilon polish with Illumina PCR-free reads brought all of the assemblies to a similar base quality level as observed by variant detection and also protein-coding gene prediction (Figure S4).

As an independent validation of our assemblies we used BioNano optical genome maps to test the assembly quality and contiguity. BioNano maps are now routinely used to identify structural genome variations [2] and scaffold sequencing contigs [18]. We have expanded their utility by using the inherent long-range physical/optical information to screen for chimeric sequencing contigs, collapsed and artificially expanded regions larger than 5 kb (Figure 3A,B,C); and also exploited them as a proxy for base-quality through the assessment of false-positive (sites that were nicked even though they are different from the recognition sequence of the restriction enzyme used) and false-negative nicking sites (sites that had the recognition sequence but were not nicked) (Figure S2). These erroneous regions occur in highly repetitive regions, as previously described for Col-0 [2], yet within long-read assemblies at a much lower rate compared to the BAC-based TAIR10 reference. Once again, BioNano maps confirmed that the base quality and contiguity of the ONT assemblies was on par with or in some cases surpassed the PB assemblies and current reference TAIR10.

Ultra-long read reference genomes present new opportunities to access biologically relevant variation that is even recalcitrant to the most trusted genomic resources. Despite having had access to Sanger-sequence BACs from the SG3i QTL region, we had been previously unable to fully resolve the complex genomic structure of this region [21]. The rapidly generated KBS-Mac-74 ONTmin assembly enabled us to resolve the SG3i region in less than 30 minutes.

In conclusion, we have demonstrated that highly contiguous genome assemblies with high per-base quality have indeed the potential to resolve functionally important presence/absence polymorphisms in *A. thaliana* accessions. By generating multiple such assemblies, the true nature of presence/absence polymorphisms across this species will come into much sharper relief.

## Methods

### Plant Growth

*Arabidopsis thaliana* accession KBS-Mac-74 (accession 1741) was grown in the greenhouse under 16 hr light and 8 hr dark at 22°C. Plants were put into complete dark for 48 hrs before DNA extraction, in order to reduce starch accumulation.

### Oxford Nanopore MinION Sequencing

5 g of flash frozen leaf tissue was ground in liquid nitrogen and extracted with 20 mL CTAB/Carlson lysis buffer (100mM Tris-HCl, 2% CTAB, 1.4M NaCl, 20mM EDTA, pH 8.0) containing 20μg/mL proteinase K for 20 minutes at 55°C. The DNA was purified by addition of 0.5x volume chloroform, which was mixed by inversion and centrifuged for 30 min at 3000 RCF, and followed by a 1x volume 1:1 phenol: [24:1 chloroform:isoamyl alcohol] extraction. The DNA was further purified by ethanol precipitation (1/10 volume 3 M sodium acetate pH 5.3, 2.5 volumes 100% ethanol) for 30 minutes on ice. The resulting pellet was washed with freshly-prepared ice-cold 70% ethanol, dried, and resuspended in 350 μL 1x TE buffer (10 mM Tris-HCl, 1 mM EDTA, pH 8.0) with 5 μL RNase A (Qiagen, Hilden) at 37°C for 30 min, followed by incubation at 4°C overnight. The RNase A was removed by double extraction with 24:1 chloroform:isoamyl alcohol, centrifuging at 22,600x*g* for 20 minutes at 4°C each time. An ethanol precipitation was performed as before for 3 hours at 4°C. The pellet was washed as before and resuspended overnight in 350 μL 1x TE.

Genomic DNA sample was further purified for ONT sequencing with the Zymo Genomic DNA Clean and Concentrator-10 column (Zymo Research, Irvine, CA). The purified DNA was then prepared for sequencing following the protocol in the genomic sequencing kit SQK-LSK108 (ONT, Oxford, UK). Briefly, approximately 1 μg of purified DNA was repaired with NEBNext FFPE Repair Mix for 60 min at 20°C. The DNA was purified with 0.5X Ampure XP beads (Beckman Coulter). The repaired DNA was End Prepped with NEBNExt Ultra II End-repair/dA tail module including 1 μl of DNA CS (ONT, Oxford, UK) and purified with 0.5X Ampure XP beads. Adapter mix (ONT, Oxford, UK) was added to the purified DNA along with Blunt/TA Ligase Master Mix (NEB, Beverly, MA) and incubated at 20°C for 30 min followed by 10 min at 65°C. Ampure XP beads and ABB wash buffer (ONT, Oxford, UK) were used to purify the library molecules and they were recovered in Elution buffer (ONT, Oxford, UK). Purified library was combined with RBF (ONT, Oxford, UK) and Library Loading Beads (ONT, Oxford, UK) and loaded onto a primed R9.4 Spot-On Flow cell (FLO-MIN106). Sequencing was performed with a MinION Mk1B sequencer running for 48 hrs. Resulting FAST5 files were base-called using the Oxford Nanopore Albacore software (v0.8.4) using parameters for FLO-MIN106, and SQK-LSK108 library type.

### Pacific Biosciences RSII Sequencing

20 g young leaf material were frozen in liquid nitrogen and ground to fine powder using mortar and pestle. The powder was directly transferred into lysis buffer and DNA was extracted with a Qiagen genomic DNA extraction kit (Qiagen, Hamburg, Germany) in combination with Qiagen genomic tip columns (500/G; Qiagen, Hamburg, Germany) according to the manufacturer’s protocol. DNA quality and quantity was determined with a NanoDrop ND 1000 spectrometer (PeqLab, Erlangen, Germany), a Qubit 2.0 Fluorometer (Thermo Fisher Scientific, Waltham, USA) and by pulse field gel electrophoresis. A total of 10 μg genomic DNA was sheared to a target fragment size of 30 kb using a Megaruptor™ 2 device (Diagenode, Denville, USA). A 20 kb template library was prepared using the BluePippin™ size-selection system according to the manufacturer’s protocol (P/N 100-286-000-07, Pacific Biosciences, California, USA). The final library was sequenced on a Pacific Biosciences RSII instrument following the Magbead loading protocol at the Max-Planck-Genome-Centre Cologne, Germany (http://mpgc.mpipz.mpg.de). A total of 18 SMRT cells were run resulting in 8.4 Gb of sequence.

### Sequence extraction, assembly, consensus and correction

Raw ONT reads (fastq) were extracted from base-called FAST5 files using poretools [23]. Raw PB reads (fastq) were extracted directly from the base-calling output of the RSII instrument. Overlaps were generated using minimap [14] with the recommended parameters (-Sw5 -L100 -m0). Genome assembly graphs (GFA) were generated using miniasm [14]. Unitig sequences were extracted from GFA files. Canu assemblies were generated using the default parameters and the complete canu pipeline [Citation error]. Three rounds of consensus correction was performed using Racon [15] based on minimap overlaps, and the resulting assembly was polished using Illumina PCR-free 2x250 bp reads mapped with bwa [24] and pilon [16]. Genome stats were generated using QUAST [25].

### Preparation of BioNano optical genome maps

High molecular weight (HMW) DNA was extracted from up to 5 g fresh leaf tissue using a modified BioNano Genomics protocol [2]. Briefly, extracted HMW DNA was nicked with the enzyme Nt.BspQI (NEB, Beverly, MA, USA), fluorescently labeled, repaired and stained overnight according to the BioNano Genomics nick-labeling protocol [26]. KBS-Mac-74 nick-labelled DNA was run on a single flow cell on the Irys platform (BioNano Genomics, San Diego, CA, USA), for 90 cycles to generate 22.5 Gb raw data. The IrysView software (BioNano Genomics; version 2.5.1) was used to quality filter the raw data (>100kb length, >2.75 signal/noise ratio) and molecules were assembled into contigs using unaltered the “human genome” parameters. Resulting BNG cmaps were compared against the different assemblies using RefAlign [2], and collapsed regions or artificial expansions were detected as structural variations using the structomeIndel.py script (https://github.com/RyanONeil/structome).

### Variant analysis, Annotation and genome comparisons

Variant analysis was based on mapping with BWA [24] and summarizing using samtools [27] and bcftools [28]. Assemblies were repeat masked using RepeatMasker [19], and genes were called with Augustus *Arabidopsis thaliana* gene models [29]. KBS-Mac-74 specific centromere (158 and 178 bp monomers) and rDNA (26S, 5.8S, 18S, and 5S) repeats were identified in the PBcan assembly with blast (blastn) using representative sequence from Col-0 TAIR10; the KBS-Mac-74 specific repeats were then used to search all four assemblies. For SG3/SG3i analysis, the complete genomic region for At4g30720 was blasted against the KBS-Mac-74 ONTmin iteration 4 assembly. 50 kb upstream and downstream of the two At4g30720 KBS-Mac-74 ONTmin hits were used to align to the Col-0 TAIR10 assembly to identify insertions, deletions and rearrangement.

## Data availability

Raw sequencing data was deposited in the European Nucleotide Archive (ENA) under project PRJEB21270. Raw BioNano Genomics molecules and assembled maps are deposited under BioProject ID PRJNA390205. Final polished assemblies were deposited in the ENA Genome Assembly Database. Code and intermediate results were deposited at GitHub (https://github.com/fbemm/onefc-oneasm).

## Contributions

TPM, FJ, FB, DW and JRE conceived the study, analyzed the data and wrote the manuscript. STM and FB generated the ONT and PB sequence respectively. FJ and JPS generated and analyzed BNG data. OL and TPM analyzed the SG3/SG3i QTL.

## Acknowledgement

We thank the Max Planck Genome Centre Cologne for performing Pacific Biosciences long read sequencing in this study. This work was funded by ERC AdG IMMUNEMESIS and the Max Planck Society. FJ was supported by a Human Frontier Science Program Organization long-term fellowship. JRE is an investigator of the Howard Hughes Medical Institute.

